# Adenylyl cyclase 9 as a molecular scaffold to dissect the regulatory mechanisms of membrane adenylyl cyclases

**DOI:** 10.64898/2026.04.18.719376

**Authors:** Ilayda Kantarci, Haoriwa, Volodymyr M. Korkhov

## Abstract

Adenylyl cyclases (ACs) convert ATP into the second messenger cAMP, thus directly influencing cellular signaling in response to a wide variety of stimuli. Despite their physiological importance, structural studies of isoform-specific AC regulation are compounded by difficulties in AC expression and purification. Here, we designed a chimeric construct AC95, combining human AC9 as a molecular scaffold and incorporating the catalytic-allosteric core of human AC5. Cryo-EM analysis of AC95 at 3.5 Å resolution revealed a state of AC95 bound to both ATPαS and forskolin, demonstrating that the chimera reproduces AC5-like allosteric regulation while retaining the structural features of the AC9 scaffold. Although AC95 chimera retained the ability to bind to and be activated by forskolin, it lost the ability to be autoinhibited by the C2b domain of AC9. Moreover, AC95 is insensitive to inhibition by specific AC5 inhibitors SQ22,536 and NKY80, suggesting that these molecules may target a site distinct from the catalytic-allosteric core of AC5 grafted into the AC95 chimeric construct. Our results establish a generalizable approach for investigating isoform-specific regulation of membrane ACs by small molecules, offering a potential path for structure-based drug discovery targeting distinct AC isoforms.

## INTRODUCTION

Adenylyl cyclases (ACs) are integral membrane enzymes that catalyze the conversion of ATP into cyclic adenosine monophosphate (cAMP), a versatile second messenger that governs diverse cellular processes, including transcription, metabolism, ion channel activity, and signal transduction [1, 2]. Nine membrane AC isoforms (AC1–AC9) have been identified in mammals, each with distinct tissue distributions, regulatory properties, and physiological roles [3]. Although all AC isoforms share a conserved catalytic domain architecture, the AC isoforms differ in their responsiveness to G proteins, Ca²⁺/calmodulin, phosphorylation, and other modulatory inputs. This enables ACs to intricately regulate cellular signaling and cellular responses in a cell type- and tissue-specific manner [3-5].

Early X-ray crystallographic studies of mammalian ACs focused on the chimeric catalytic domains isolated from AC5 and AC2 in complex with a small molecule AC activator, forskolin, nucleotide analogues and the stimulatory subunit of the heterotrimeric G protein (Gαs) [6]. These studies established the two-metal ion mechanism of ATP-to-cAMP conversion, revealed intermediate catalytic states, and provided a plausible explanation for forskolin-mediated AC activation [7-9]. Although forskolin is a non-physiological activator and no endogenous small molecule ligand for its binding site in the ACs (the allosteric site) has been identified to date, these experiments demonstrated that both Gαs and forskolin are required to stabilize the C1a–C2a catalytic dimer [6, 10-12]. However, full-length membrane AC structures remained unresolved for over two decades, owing to the challenges of expressing and purifying ACs as membrane proteins produced only in limited amounts [13].

The first cryo-EM structure of a full-length AC, the bovine AC9 (bAC9) bound to Gαs, revealed the 12-helix transmembrane (TM) bundle, the coiled-coil organization of the extended helices connecting the catalytic domains C1a and C2a to the TM6 and TM12, respectively [13]. This structure also captured Gαs binding and an autoinhibited, or occluded, state of the AC, where a portion of the C-terminal C2b domain is inserted directly into the active and allosteric pocket of bAC9, thus blocking the catalytic activity [13, 14]. Deletion of C2b relieved autoinhibition and allowed forskolin and a nucleotide analogue, MANT-GTP, to bind to the allosteric and active site of the enzyme, respectively [13]. A recent cryo-EM structure of human AC9 (hAC9) confirmed these observations, providing further insights into [15].

The regulatory properties of AC9 make it unique among AC isoforms [16]. It is expressed in a wide range of tissues, playing roles in regulating cardiac contractility, smooth muscle tone, and neuronal signaling [5, 16, 17]. AC9 dysregulation has been implicated in cardiovascular and metabolic disorders, underscoring its physiological importance and therapeutic potential [18, 19]. Beyond these functional roles, AC9 has emerged as a benchmark for mammalian AC research, providing a powerful platform for structural and mechanistic investigations that illuminate both the general principles of AC regulation and the isoform-specific features that distinguish individual family members [13, 15, 20]. AC9 differs from other AC isoforms in that Gαs to date is the only known natural activator, with the C2b domain playing an autoregulatory role [13, 14, 16]. Although traditionally considered forskolin-insensitive, AC9 can be activated by forskolin at high concentrations in the presence of Gαs [13], highlighting its complex regulatory properties.

The highly dynamic nature of membrane ACs continues to complicate detailed mechanistic studies of these enzymes. For example, a recent cryo-EM study of hAC5 produced a 7-Å structure of the hAC5–Gβγ complex, revealing the Gβγ interaction interface, but failing to resolve the catalytic domain of hAC5, which appeared to be disordered or flexible [21]. Similarly, a structure of the dimeric species of hAC5 at low resolution showed the remarkable potential of the structural studies to shed light on the mechanisms of AC complex assembly and regulation [21]. However, the flexibility of the protein precluded obtaining the structure at high resolution. Similarly, our recent study of the structure and regulation of bAC8 and its complexes with Gαs, Gβγ and calmodulin provided only limited insights into the molecular determinants of bAC8 modulation by these agents. The structure of bAC8-Gαs in the presence of calmodulin revealed only the ordered part of the complex that included only parts of bAC8-Gαs [22]. Although we were able to map the interaction between bAC8 and Gβγ using mass spectrometry-based analysis, the complexity of the Gβγ-bound bAC8 complex limited our ability to obtain a cryo-EM-based 3D reconstruction [22]. Furthermore, several specific small molecule membrane AC inhibitors have been identified, and their mode of action via the AC catalytic site has been postulated [23, 24]. These compounds hold promise as potential AC-targeted therapeutics, which could provide a starting point for further rational design of new AC modulators. However, to this end there is no direct experimental validation of the structural basis and mode of action of these small molecules. These examples highlight the need for developing new approaches to study membrane ACs.

Here, we analyzed the cryo-EM structures of hAC9 bound to Gαs in the presence of ATPαS and forskolin, revealing important new features of hAC9-Gαs complex complementing the previously determined structures of bAC9 [13, 20] and hAC9 [15]. Our analysis revealed that the purified hAC9-Gαs complex appears to exist as an ensemble of conformations, including two distinct states: the occluded state and the ATPαS–bound state. Building on the experimental cryo-EM structures of hAC9 and on computational structure predictions, we developed a chimeric construct integrating the catalytic-allosteric core of hAC5 into the catalytic domain of the hAC9 scaffold. The experiments utilizing this construct provide important insights into AC regulation, validating the use of such chimeric constructs as a new general approach to studying membrane ACs as dynamic enzymes and potential targets for small-molecule therapeutics.

## RESULTS

### Conformational ensembles of hAC9 bound to Gαs protein

hAC9 consists of two six-helix transmembrane domains (TM1–6 and TM7–12) connected to cytosolic catalytic domains (C1a and C2a) that assemble to form the catalytic core (Figure 1A-C). To gain insights into the structural basis of hAC9 activation, we determined cryo-EM structures of hAC9 in complex with Gαs, both in the absence and presence of ATPαS and forskolin. Comparisons with recently published Gαs-bound and apo hAC9 structures [15] (Figure 1D, E) shows that Gαs binding stabilizes a single overall conformation containing the C2b domain. However, our 3D variability analysis, used to address the intrinsic flexibility of hAC9, revealed the presence of two distinct conformational states (Supplementary Figure S2). In the presence of ATPαS and forskolin, the hAC9-Gαs complex adopts two distinct conformations (Figure 1F, G) (Supplementary Figure S2), in contrast to the single occluded conformation observed in the absence of nucleotide and forskolin (Supplementary Figure S3). In the first conformation, a fragment of the C2b domain folds into the catalytic core, occluding the active site (Figure 1F). In the second conformation, ATPαS is bound at the catalytic site, and the C2b domain is displaced such that it no longer occludes the catalytic center (Figure 1G). Comparison with the recently published apo structure shows that Gαs binding displaces the α4 helix (Cα RMSD ≈ 4 Å for residues 512–522) (Figure 1H-K), allowing the C2b domain (residues 1260-1277) to fold into the catalytic–allosteric core. In the occluded state, the C2b domain blocks substrate access through stabilizing hydrogen bonds between Q522–R1263 and N515–V1266 (Figure 1J). The occluded conformation is similar to that previously reported for bAC9-Gαs [13] and in hAC9-Gαs [15] (Supplementary Figure S9). In the nucleotide-bound conformation the α4 helix undergoes further outward displacement, partially relieving C2b-mediated occlusion and creating space for ATPαS binding (Figure 1K).

**Figure 1:**
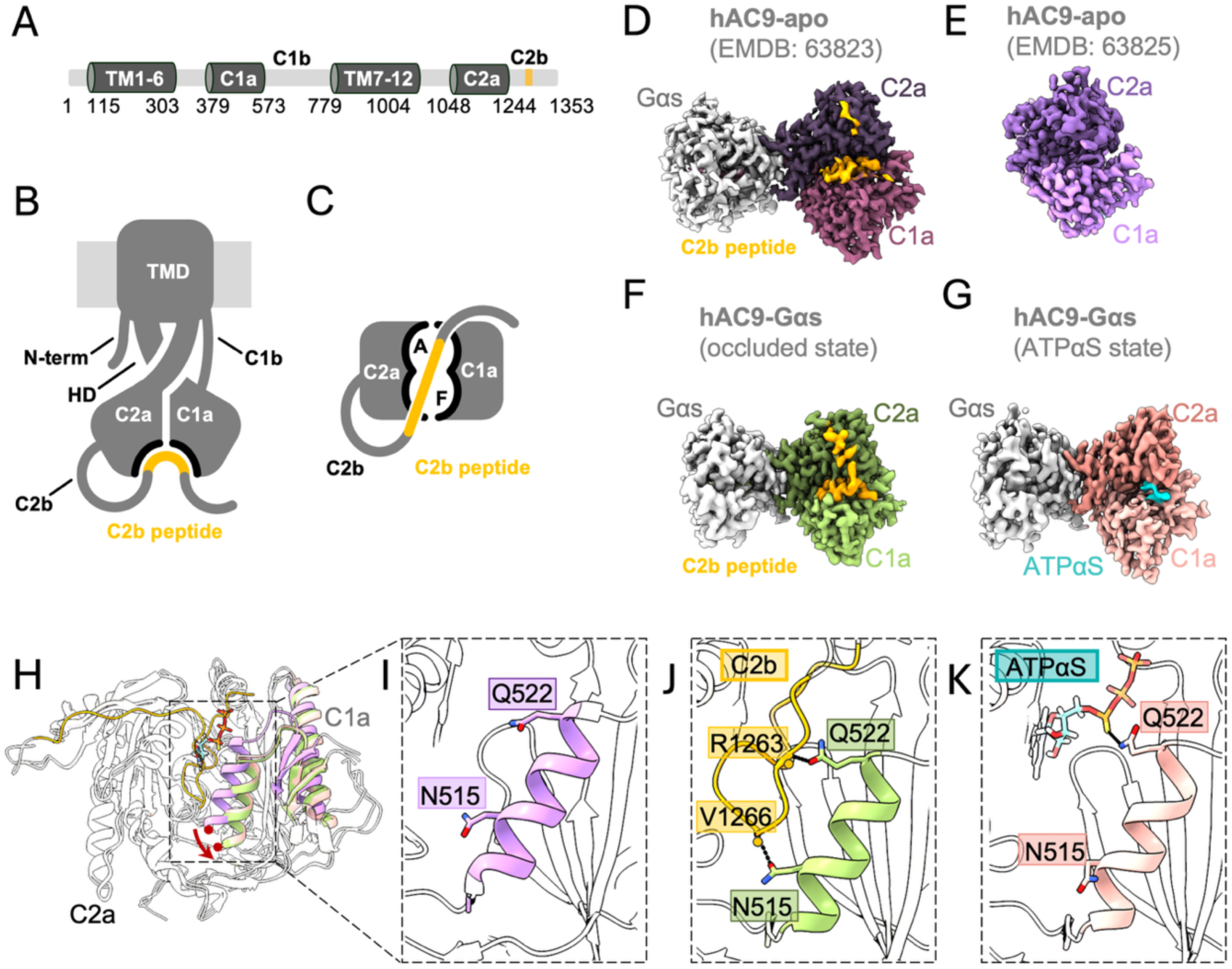
Conformational dynamics of hAC9. **A-C.** Overall architecture of human AC9 (hAC9), highlighting the allosteric, forskolin binding **(C)**, and catalytic, ATP binding, sites and occluded by the C2b peptide. **D.** Cryo-EM maps of hAC9-Gαs, EMDB: 63823, **E.** hAC9, EMDB: 63825, F. hAC9–Gαs–ATPαS–forskolin complex (green) showing the C2b domain (orange) occluding the catalytic center, occluded state **G.** hAC9–Gαs–ATPαS–forskolin, with ATPαS (cyan) bound to the catalytic site; no density corresponding to forskolin is observed, ATPαS-bound state. **H.** Structural comparison of hAC9, with the C2a domain as the reference, reveals a relative movement of the α4 helix upon Gαs binding. **I-K.** Rearrangement of residues within the α4 helix coordinates both C2b-mediated occlusion and ATPαS binding.

### Forskolin is excluded from its binding site in the nucleotide-bound state of AC9-Gαs

Despite including forskolin during sample preparation, we did not observe a clearly defined density corresponding to forskolin in the allosteric binding pocket of the ATPαS-bound hAC9-Gαs structure (Figure 1G). This suggests that the ATPαS-bound state of hAC9 captured here adopts a conformation that does not favor forskolin binding, indicating that additional conformational changes may be required to fully accommodate this allosteric activator. These findings underscore the state-dependent regulation of hAC9, with Gαs and nucleotide binding driving distinct structural transitions that modulate active site accessibility and isoform-specific catalytic activity.

### Rational design AC9-based chimeric AC5 constructs b9AC5, AC95

Chimeric construct-based structural studies combining the C1a domain of canine AC5 with the C2a domain of rat AC2, provided important structural insights into the catalytic mechanism of adenylyl cyclases [6, 23, 25-27], but they were limited in explaining isoform-specific catalytic and allosteric regulation. Our previous cryo-EM studies of the bAC9 resolved multiple conformational states, revealing key aspects of its catalytic activation and regulation [13, 20]. Combined with the current study and the recent structure of hAC9 [15], a more complete picture of the molecular features of AC9 is emerging. However, a major challenge remains in structural studies of membrane AC isoforms other than AC9: to date only the structure of one other mammalian full-length AC at high resolution could be determined, the bAC8 [22]. All other mammalian ACs currently remain inaccessible to direct structural studies, largely due to challenges in expression, purification, stability and conformational heterogeneity. This is illustrated by the recent work on hAC5, which revealed a largely unresolved catalytic domain [21].

To address these challenges and enable structure–function studies of membrane AC isoforms, we designed two complementary chimeric strategies based on AC9. To benchmark these two approaches, we chose hAC5 as a test case because of its biomedical significance and the unresolved catalytic domains in recently published cryo-EM analysis [5, 21, 28]. In the first approach, the entire catalytic core (C1a and C2a domains) together with the part of the helical domain of hAC5 was incorporated into the transmembrane scaffold of bAC9, generating a construct named “b9AC5” (Figure 2A-D). The design of this chimera was guided by structural predictions of hAC5 together with the available cryo-EM structure of bAC9 (Figure 2E). Structural predictions of the b9AC5 construct showed that the helical and catalytic domains closely resemble those of hAC5 and display comparable pLDDT confidence values, supporting the structural integrity of the designed chimera (Figure 2F). The b9AC5 construct was subsequently expressed and purified. However, size-exclusion chromatography (SEC) revealed a broad elution profile, and SDS–PAGE analysis did not show a clear band at the expected molecular weight (Supplementary Figure S1 E, F), indicating sample heterogeneity. Consistent with this observation, cryo-EM analysis of the construct resulted in reconstructions with limited resolution, preventing reliable model building and detailed structural interpretation (Figure 2G) (Supplementary Figure S5).

**Figure 2:**
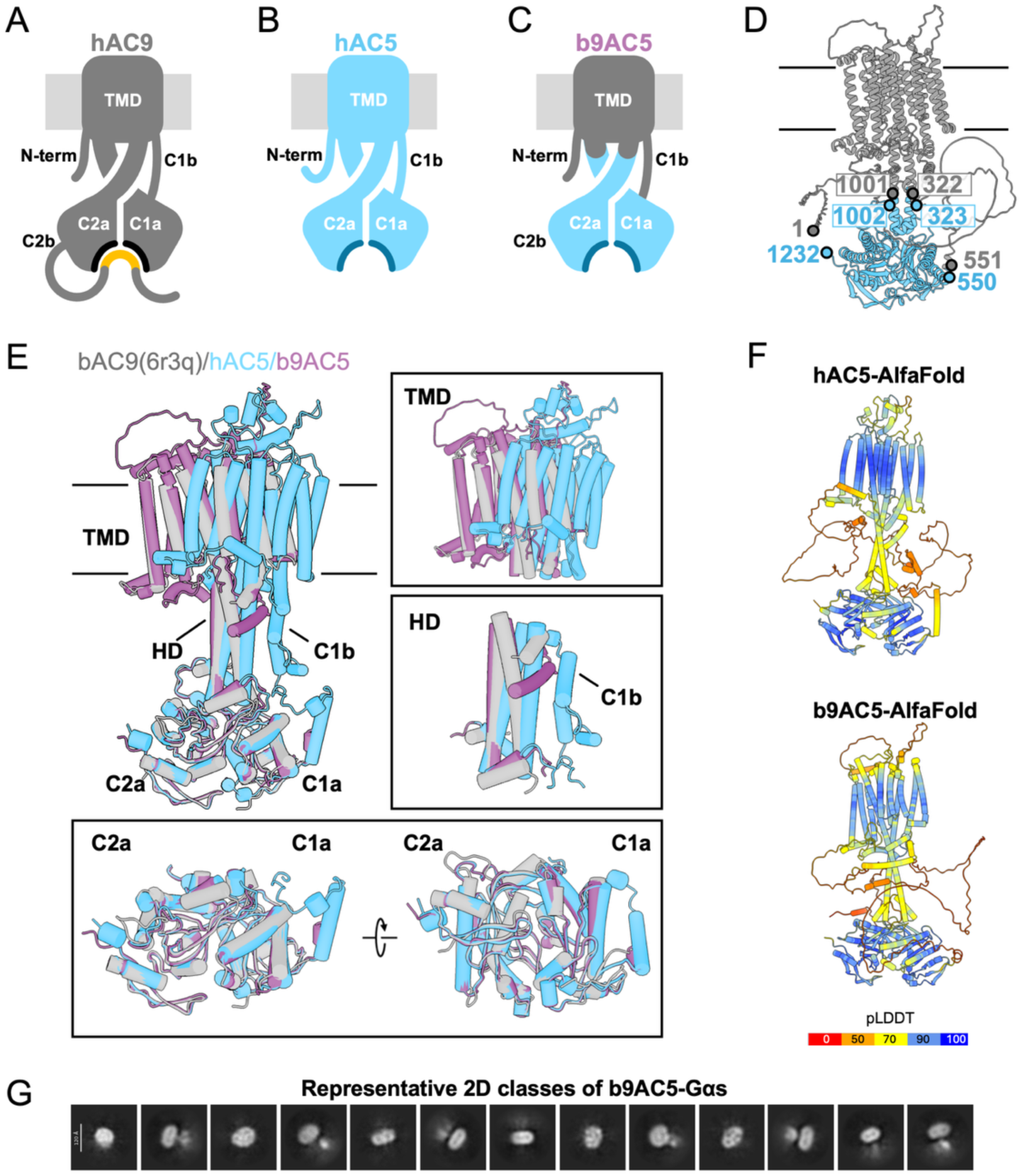
b9AC95 chimera design. **A-C.** Schematic representation of **(A)** bAC9, **(B)** hAC5, and **(C)** b9AC5, **D.** Residues in b9AC5, 1–322 and 551–1001, derived from bAC9, form the anchor (TMD, helical domain, C1b), whereas residues 323–550 and 1002–1232, derived from hAC5 (C1a–C2a), constitute the catalytic domains. **E.** Structural comparison of bAC9, and AlfaFold3 predictions of hAC5, and b9AC5 were guided the design. F. AlphaFold**3** predictions of hAC5 and b9AC5. **G.** Cryo-EM data analysis of b9AC5 showed AC-like particles in 2D classification, but clear structural features of the catalytic domains were not observed.

To overcome these limitations and improve structural stability and particle homogeneity, we pursued an alternative chimeric strategy in which hAC9 served as the structural scaffold, while the catalytic and allosteric core of hAC5 was incorporated (Figure 3A-D). Sequence alignments (Figure 3E), structural comparisons with early crystallographic models, and our hAC9 structures guided the design to maintain structural compatibility (Figure 3F-H). The resulting chimeric construct featuring 11 amino acid substitutions, which we refer to here as “AC95”.

**Figure 3:**
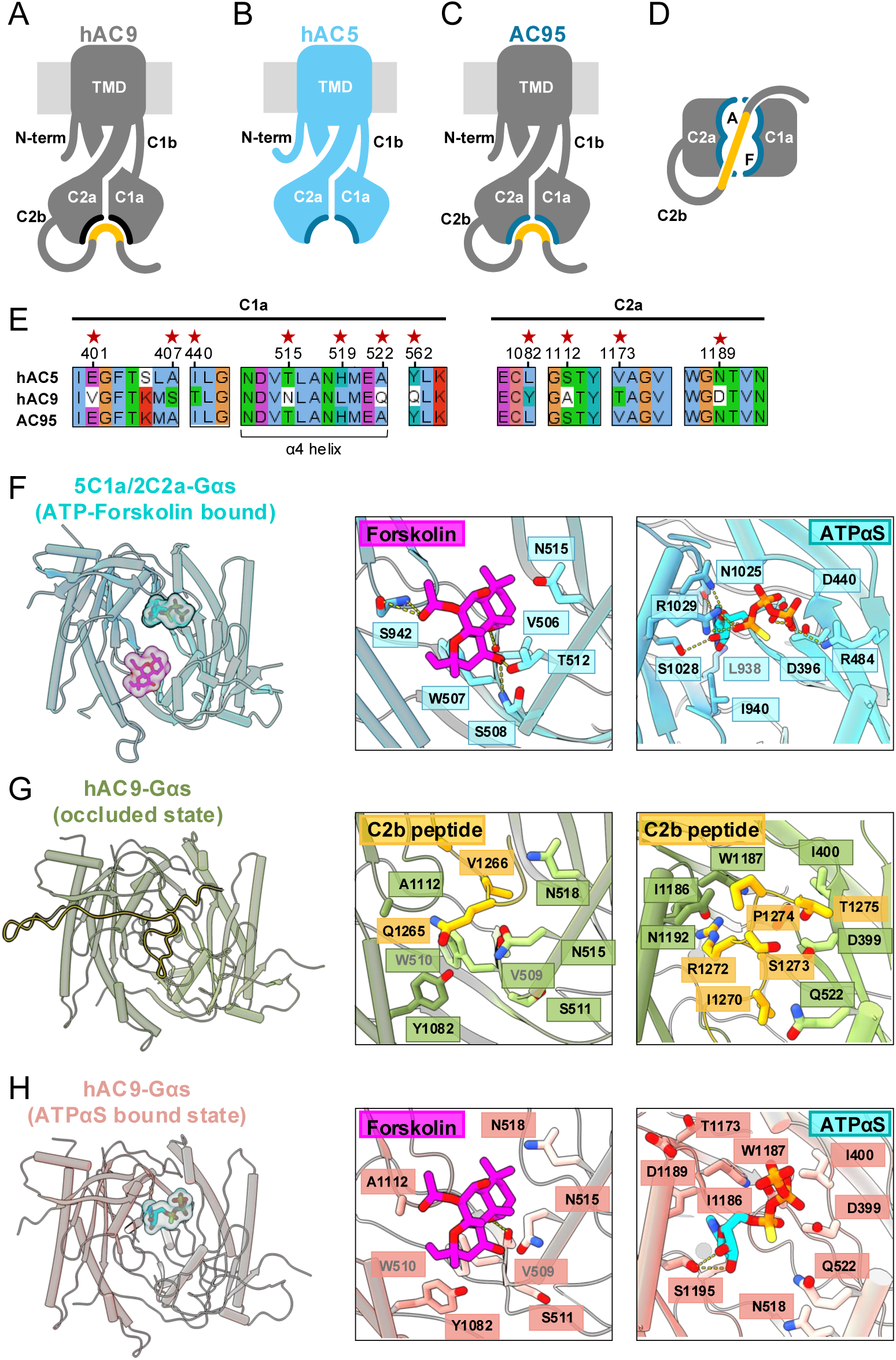
AC95 chimera design. A-D. Schematic representation of (A) hAC9, (B) hAC5, and (C-D) AC95, where AC9 retains its regulatory and membrane-proximal elements while incorporating the catalytic core of hAC5. E. Sequence alignments of hAC9, hAC5, and AC95 and the red star indicated the swapped residues for AC95 (V401E, S407A, T440I, N515T, L519H, Q522A, Q562Y, Y1082L, A1112S, T1173V, D1189N). F. Crystallographic models of the catalytic and allosteric sites based on the 5C1a/2C2a structure, with ATPαS docked into the active site. G-H. Residues within ∼3.5 Å of the catalytic and allosteric sites in the hAC9–Gαs occluded (G) and ATPαS-bound (H) states were analysed, including those surrounding the forskolin-binding pocket by docking the forskolin to its binding site. Residues in proximity (3.5 Å) in AC9 catalytic site were subsequently replaced by their AC5 counterparts.

### Biochemical validation of the AC95 construct

To determine suitability of AC95 for experimental analysis of structure and function, we expressed the protein in mammalian cells and purified it using a procedure used for hAC9 (see Materials and Methods). Size-exclusion chromatography revealed a predominantly monodisperse population comparable to hAC9, and SDS–PAGE was confirmed the expected molecular weight and purity of the protein (Supplementary Figure 1).

Next, we investigated the basal and stimulated AC activities of the AC95 chimera and compared it to the purified wild type hAC9 and hAC5 using a standard AC activity assay in the presence of allosteric site activators and catalytic site inhibitors (Figure 4A, B). AC95 showed lower basal activity than both hAC9 and hAC5 (Figure 4, C-E). Despite this reduction, AC95 showed robust response to the activated Gαs protein (Figure 4E).

**Figure 4:**
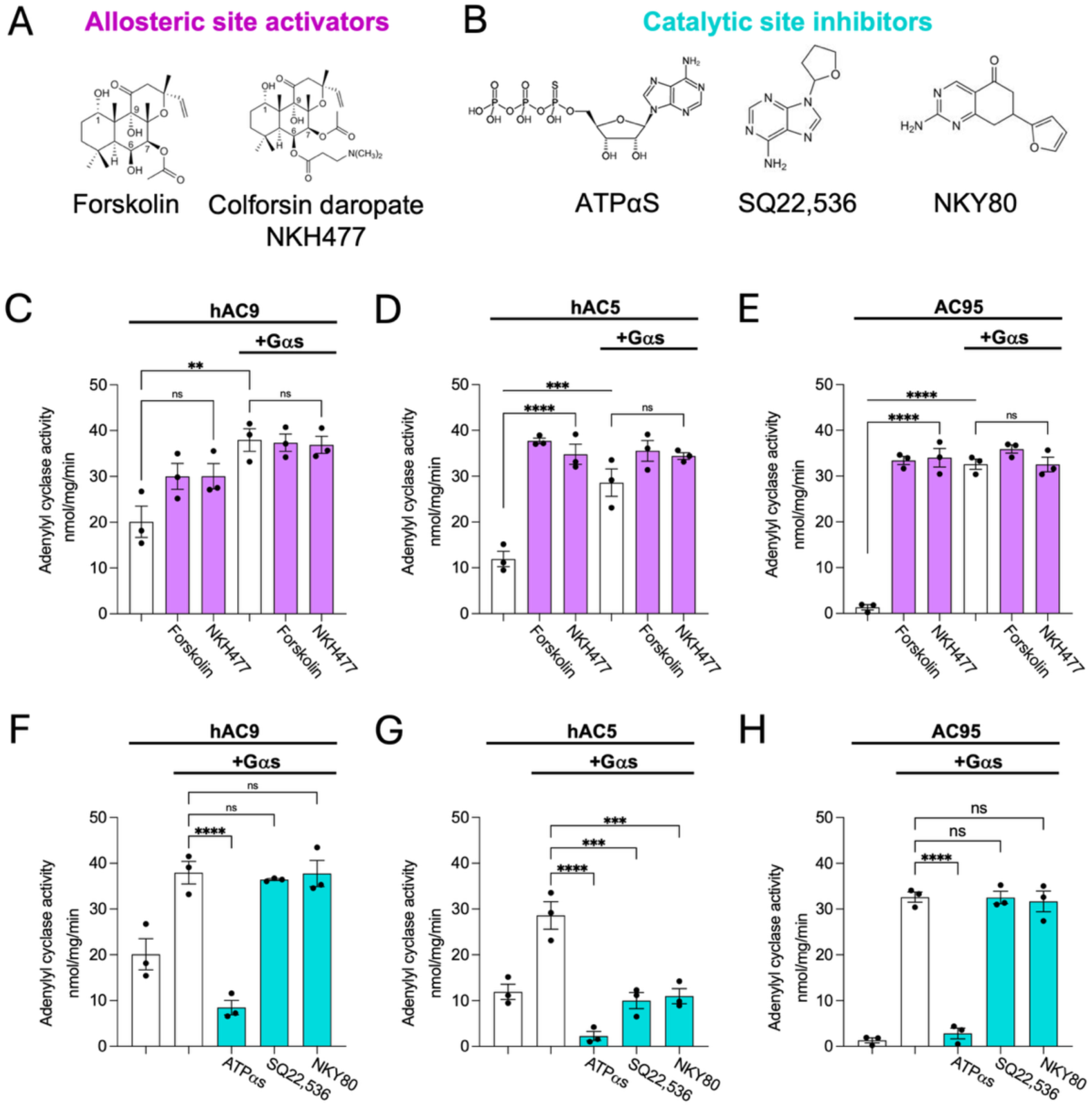
Biochemical analysis of hAC9, hAC5, and AC95. **A.** Allosteric-site activators **B.** catalytic-site inhibitors used in this study. **C-E.** Enzymatic activity of hAC9, hAC5, and AC95 in the presence of Gαs, forskolin, and NKH477 **F-H.** Enzymatic activity of hAC9, hAC5, and AC95 in the presence of Gαs, ATPαS, SQ22,536, and NKY80. For all experiments, the data are shown as mean ± standard error of the mean (SEM) (*n* = 3). Statistical significance was assessed using one-way ANOVA, followed by Tukey’s multiple comparisons test, with *P* < 0.0001 (“****”), *P* = 0.0006 (“***”), and *P* = 0.0385 (“*”). No significant difference is indicated as “ns”.

We also tested the response of AC95 to forskolin and the water-soluble forskolin analog NKH477[23, 29]. AC95 showed an enhanced response to both activators compared to the wild type hAC9 (Figure 4C, E). These results suggest that the catalytic-allosteric core of hAC5 confers properties that enhance small molecule responsiveness in the chimeric enzyme (Figure 4D, E).

The wild-type hAC9 was only insignificantly stimulated by forskolin and showed an approximately 2-fold activation by Gαs (Figure 4C). In contrast, hAC5 showed a more pronounced significant forskolin and Gαs-mediated activation, with each treatment individually producing maximal stimulation of catalytic activity (Figure 4D). Maximal AC95 activity was reached upon addition of either Gαs or forskolin individually, indicating that each activating agent maximally stimulated cAMP production (Figure 4E). These results demonstrate that AC95 is efficiently expressed, biochemically active, and responsive to the classical activators of AC activity (forskolin and NKH477).

### AC95 is inhibited by ATPαS but not SQ22,536 and NKY80

To determine whether grafting the hAC5 catalytic-allosteric core into hAC9 confers the sensitivity of the resulting construct to the AC inhibitors, we tested SQ22,536 [23]and NKY80 [23]in our activity assays. These compounds were previously suggested to act as membrane-permeable nucleotide analogs capable of inhibiting AC5 and AC6 [23, 30, 31], presumably blocking the enzymatic activity by binding to the active site of membrane ACs. The positive control, ATPαS, a “non-cyclizable” ATP analog, effectively inhibited AC activity under all tested conditions (Figure 4, F-H) consistent with its action as a competitive ATP site inhibitor.

In contrast, SQ22,536 and NKY80 failed to inhibit both hAC9 and AC95 activity in our assay (Figure 4F, H). Interestingly, these compounds did inhibit Gαs-activated hAC5 (Figure 4G) as expected, highlighting a differential sensitivity between the chimeric enzyme and hAC5 (Figure 4G, H). The observed lack of inhibition of AC95 by SQ22,536 and NKY80 (Figure 4H) may indicate that these compounds may not interact directly with the hAC5-derived catalytic core when embedded in the hAC9 scaffold, or that they may bind to an alternative regulatory site (or sites) present in hAC5 but are absent or inaccessible in AC95 and in the wild-type hAC9.

### Structure of AC95 chimera bound to Gαs, forskolin, and ATPαS

To gain insights into the allosteric regulation of the AC95 chimera, we prepared cryo-EM samples and determined its structure in complex with Gαs-GTPγS, forskolin, and ATPαS using single-particle analysis, achieving an overall resolution of 3.8 Å (Figure 5A) (Supplementary Figures S4,6-7 Table S1). The structure revealed all the canonical features previously observed in hAC9, confirming that the overall architecture of AC95 is largely preserved despite the chimeric replacement of the catalytic core in the hAC9 scaffold. Focused local refinement of the C1a–C2a catalytic domains improved the local resolution of the cryo-EM density map to 3.5 Å (Figure 5B, Supplementary Figure S4,6), enabling visualization of bound ATPαS and forskolin within the active and allosteric sites, respectively (Figure 5B).

**Figure 5:**
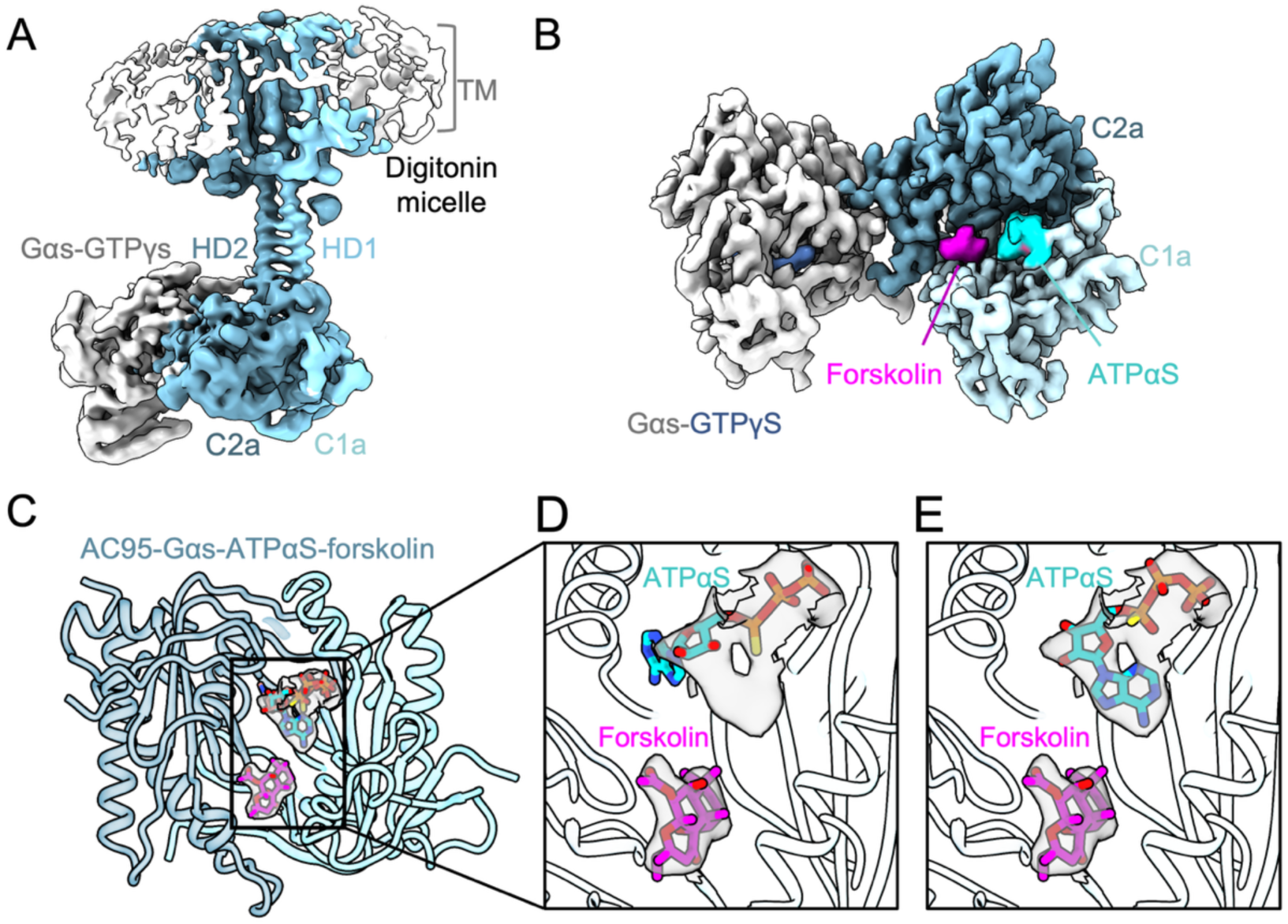
Cryo-EM map of the AC95 chimera bound to Gαs, forskolin, and ATPαS. **A.** Cryo-EM map of AC95 in complex with Gαs, forskolin, and ATPαS, showing the twelve transmembrane helices (TM), helical domain (HD), and catalytic domains C1a and C2a. **C.** The focused map of the catalytic core (C1a–C2a) reveals well-defined densities for both ATPαS and forskolin. **C.** Densities observed in the AC95 catalytic core correspond to forskolin and ATPαS. **D-E**. Two alternative ATPαS poses are fitted into densities observed in the AC95 catalytic core. **(D)** ATPαS pose 1 observed in hAC9-Gαs, fitted into the corresponding density in **(E)** AC95 ATPαS pose 2 fitted into the corresponding density in AC95

### Occluded state is not observed in AC95 chimera

Although the C2b domain is present in the AC95 chimeric protein construct, we did not observe the occluded conformation (Figure 5C) (Supplementary Figure S4). Single-particle analysis of AC95 consistently produced 3D classes of the active, Gαs-bound conformation, without the density features of the occluded state (Figure 5A, B) (Supplementary Figure S4). This contrasts with prior observations in hAC9, where the C2b domain stabilizes the occluded conformation under similar conditions (Figure 1F) (Supplementary Figure S2). Replacing the catalytic core with the hAC5-derived segment in AC95 appears to change the enzyme’s conformational landscape, reducing the tendency of the catalytic domains to adopt the occluded state, even with the regulatory C2b domain present.

### ATPαS and forskolin binding in the AC95 chimera

The densities corresponding to forskolin and ATPαS were well-defined, allowing precise assignment of their positions in the catalytic core (Figure 5C). Interestingly, the density corresponding to ATPαS binding differed from that observed in hAC9, indicating a distinct nucleotide-binding mode in the chimera (Figure 5D, E) (Supplementary Figure S10).

Comparison of the catalytic sites between the cryo-EM structures of hAC9–Gαs–ATPαS-forskolin complex and AC95–Gαs–ATPαS–forskolin complex revealed notable structural differences. In AC95, forskolin binding displaces the α4 helix outward (Cα RMSD ≈ 3.5 Å) (Figure 6A), accompanied by a distinct shift of ATPαS into pose 2, whereas wild type hAC9 predominantly exhibits pose 1 (Figure 6 A-C) (Supplementary Figure S10). This indicates that forskolin might induces an alternative nucleotide configuration relative to the catalytic residues. This is reminiscent of the effect observed in soluble AC (sAC), where binding of the allosteric inhibitor LRE1 near the α4 helix displaces the substrate analog ApCpp from a catalytically unfavorable orientation (Figure 6D–F) [31]. In sAC, this repositioning primarily involves the side-chain residue R176, whereas in transmembrane ACs, the outward movement of the entire α4 helix reshapes the catalytic core. These observations suggest that α4-mediated allosteric communication between regulatory elements and the catalytic site is conserved across AC isoforms, but the structural implementation differs, which may reflect isoform-specific differences in allosteric communication between the catalytic core, the mode of ATP binding, and the regulatory domains in different AC isoforms.

**Figure 6:**
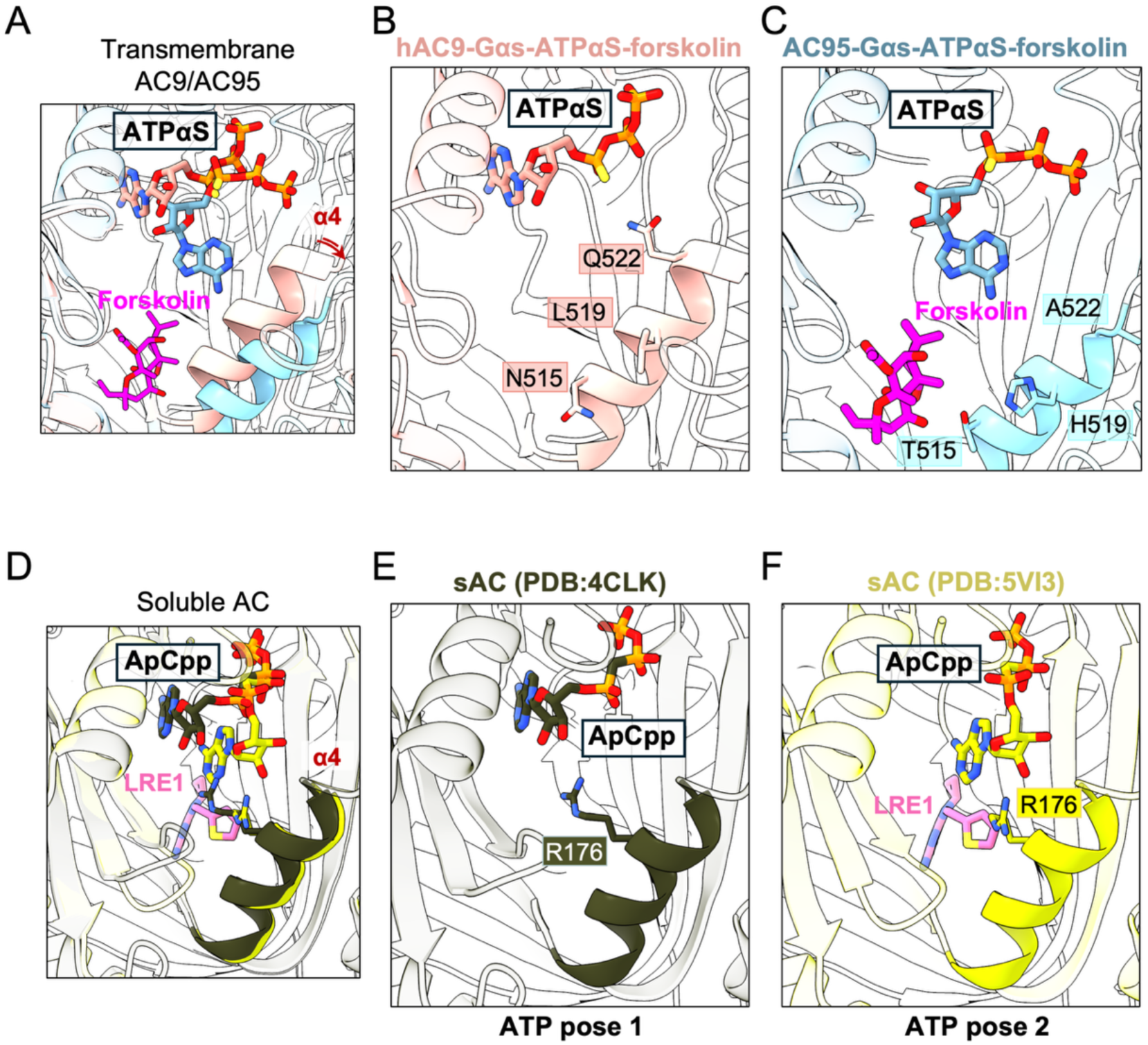
Forskolin binding alters ATPαS orientation in the AC95 chimera. **A.** Comparison of the catalytic sites of hAC9–Gαs–ATPαs-forskolin and AC95–Gαs–ATPαs–forskolin. **B-C.** In the presence of forskolin, the α4 helix is displaced outward (**B**) hAC9, the α4 helix adopts a conformation that stabilizes ATPαs in pose 1, whereas in (**C**) AC95, outward displacement of the α4 helix is associated with adoption of an alternative ATPαS binding mode (pose 2). **D-F.** In soluble AC, different binding modes of the ATP analog ApCpp are observed in the presence of the allosteric inhibitor LRE1. (**E**) In the absence of LRE1, ApCpp adopts pose 1, coordinated by residue R176 on the α4 helix. (**F**) Upon binding of LRE1 to the allosteric site, ApCpp adopts an alternative binding pose.

## DISCUSSION

The cryo-EM structures of Gαs-bound hAC9 and the chimeric AC95 highlight a coordinated interplay between the catalytic-allosteric core of the AC, the α4 helix, and the C2b domain. In the Gαs-bound state, the C2b domain occludes the catalytic core, preventing substrate access as observed in bAC9 and recent hAC9 structures. Upon nucleotide binding, the α4 helix shifts outward, partially relieving C2b-mediated occlusion and creating space for ATPαS. Despite the presence of forskolin during sample preparation, we did not observe density corresponding to this allosteric activator, indicating that the captured nucleotide-bound conformation does not favor forskolin binding (Figure 6,7). This underscores the state-dependent nature of AC9 regulation and suggests that additional conformational transitions may be required to fully accommodate allosteric ligands.

**Figure 7:**
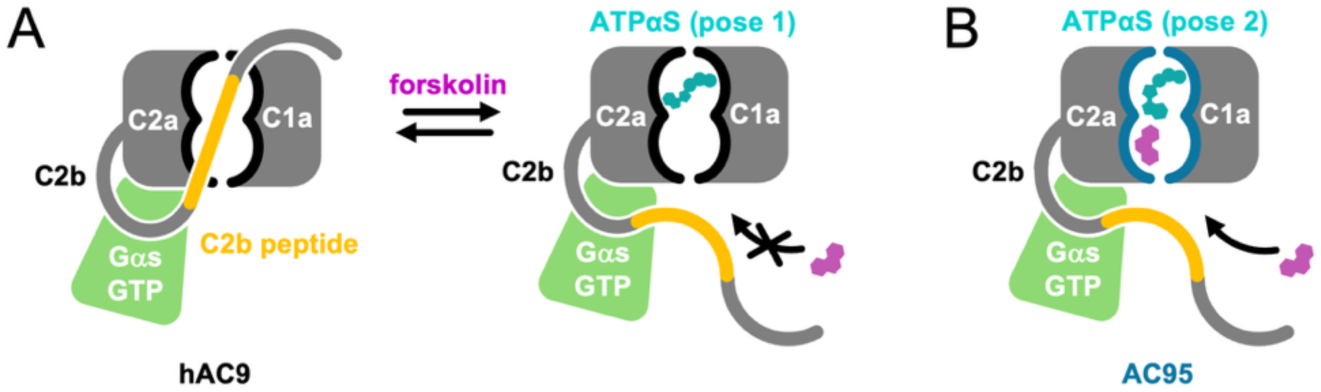
Overview of the conformational landscape of hAC9 and AC95. Based on cryo-EM structures determined in this study, hAC9 can adopt multiple conformations in the presence of an ATP analog, ATPαS, and forskolin. **A.** In the occluded state, Gαs induces the C2b domain to fold into the catalytic site, whereas in the ATPαS-bound state, Gαs stabilizes a catalytically competent conformation. **B.** In contrast, the chimeric construct featuring the hAC5-derived catalytic and allosteric core, AC95, adopts a single conformation with ATPαS and forskolin bound. The nucleotide pose is distinct from the catalysis-compatible ATP pose observed in the forskolin-free hAC9. The ATPαS pose in AC95 likely reflects a stable conformation of the forskolin-bound enzyme in which the orientation of the nucleotide analogue bound to the ATP site is not compatible with the ATP to cAMP catalytic reaction.

Incorporation of the hAC5 catalytic domain into the hAC9 scaffold preserves overall architecture but prevents the occluded state observed in wild type hAC9, allowing simultaneous binding of ATPαS and forskolin (Figure 6). Forskolin binding displaces the α4 helix outward, repositioning ATP in the active site, similar to the α4 helix-mediated ATP repositioning observed in the presence of the allosteric inhibitor LRE1 in soluble AC[31]. Analogous mechanisms have been described in other systems. In photoactivatable adenylyl cyclase (OaPAC) (Supplementary Figure S11), ATP binds in an energetically unfavorable conformation in the dark, with the AC domains remaining closed. Light activation induces structural changes only in the presence of ATP, allowing it to adopt a productive orientation for ATP to cAMP catalytic reaction [32]. Similarly, our cryo-EM reconstruction of bAC8-Gαs showed a well-resolved forskolin density and a relatively poorly defined nucleotide density in the active site [22]. These observation in different AC isoforms highlight a distinction between the transient catalytically competent nucleotide-bound states, and the stable low energy intermediates that can be resolved by structural biology techniques. In the case of AC95, the observed ATPαS pose is not conducive to cyclization, but it likely represents a previously undescribed mode of protein-nucleotide interaction in which the molecule of ATPαS is sufficiently ordered to be resolved by cryo-EM.

The chimeric AC approach can be instrumental in delineating the sites of drug action, following the approach we used here for SQ22,536 and NKY80, the membrane-permeable AC inhibitors [23, 30]. These small molecules failed to inhibit AC95 and hAC9, suggesting that (i) either the hAC9 scaffold restricts access or alters the binding properties of these compounds, or (ii) these compounds bind to hAC5 elsewhere, at a site distinct from the catalytic-allosteric core. Careful future experimentation will be required to delineate precisely the location of the small molecule inhibitor-AC interface, and chimeric construct-based methods building on the results described here may present a viable approach to solving this problem.

The utility of AC95 as a construct combining a grafted catalytic core of hAC5 within the hAC9 scaffold shows great promise in guiding future drug discovery efforts targeting membrane ACs. Remarkably, introduction of 11 mutations within the hAC9 catalytic core to produce AC95 did not compromise the biochemical properties of the protein. This hints at a possibility that this approach may be extended to cover a greater surface of the AC catalytic / regulatory sites, or to include ACs other than AC5. To this end, our attempts to express and purify full-length membrane ACs other than AC9 and AC8 for biochemical and structural studies have been largely unproductive: these proteins are biochemically unstable, are not expressed in high yields, and cannot be efficiently purified. The approach described here can now allow us to capitalize on the generally conserved membrane AC architecture and to transplant the catalytic and/or regulatory sites of interest on the surface of AC9 as a stable scaffold, enabling structure-function studies of membrane ACs that were previously inaccessible.

## Acknowledgments

This work was supported by Swiss National Science Foundation grants to V.M.K. (184951, 10003696).

## Author contributions

Conceptualization: V.M.K. Methodology: I.K., H.H., V.M.K.

Investigation: I.K., H.H., V.M.K.

Visualization: I.K., H.H., V.M.K.

Funding acquisition: V.M.K.

Project administration: V.M.K.

Supervision: V.M.K.

Writing – original draft: I.K., V.M.K.

Writing – review & editing: I.K., H.H., V.M.K.

## Competing interests

The authors declare no competing interests.

## Data and materials availability

Cryo-EM maps and models will be deposited in the Electron Microscopy Data Bank and Protein Data Bank. All other data associated with this work is available in the Source Data.

## Supplementary Materials

Materials and Methods Fig. S1 to S11 Supplementary Table 1

## Notes

### Competing Interest Statement

The authors have declared no competing interest.

