## Supplementary Information for "Adenylyl cyclase 9 as a molecular scaffold to dissect the regulatory mechanisms of membrane adenylyl cyclases"

### MATERIALS AND METHODS

**Expression of hAC9, hAC5 and chimeric AC95**. Genes encoding full-length human adenylyl cyclase 9 (hAC9; UniProt O60503), adenylyl cyclase 5 (hAC5; UniProt O95622), a chimeric b9AC5 construct (residues 1–322 and 551–1001derived from bAC9 provide the anchor (TMD, helical domain, C1b), while residues 323–550 and 1002–1232 derived from hAC5 (C1a–C2a) form the catalytic domains) and a chimeric AC95 construct (AC95; V401E, S407A, T440I, N515T, L519H, Q522A, Q562Y, Y1082L, A1112S, T1173V, D1189N), each fused to a C-terminal 3C protease site followed by an EYFP–TwinStrep tag, were synthesized and cloned by Genewiz (Azenta Life Sciences). HEK293F cells used for protein expression were maintained in suspension in FreeStyle medium supplemented with 2% fetal calf serum and 1% penicillin–streptomycin at 37 °C and 5% $CO_2$ with constant agitation. Cells were transiently transfected at a density of $2 \times 10^6$ cells $mL^{-1}$ using linear polyethyleneimine (PEI MAX; Polysciences) at a DNA:PEI ratio of 1:3 (w/w). Cells were harvested 48 h post-transfection by centrifugation, flash-frozen, and stored at −80 °C until use.

**Purification of human AC9, human AC5 and chimeric AC constructs.** HEK293F cell pellets expressing hAC9, hAC5, b9AC5 or AC95 were resuspended in buffer (50 mM Tris-HCl, pH 8.0, 150 mM NaCl) supplemented with protease inhibitors (1 mM benzamidine, 1 μg/ml leupeptin, 1 μg/ml aprotinin, 1 μg/ml pepstatin, 1 μg/ml trypsin inhibitor, and 1 μM PMSF). Cells were lysed using a Dounce homogenizer, and membranes were isolated by ultracentrifugation (Ti45 rotor, 35,000 rpm, 40 min). The membranes were solubilized at 4°C for 1 h in 1% DDM and 0.2% cholesteryl hemisuccinate (CHS), followed by ultracentrifugation under the same conditions to clarify the lysate. The resulting supernatant was incubated with CNBr-Sepharose beads coupled to an anti-GFP nanobody for 1 h. The resin was collected in a gravity column (Bio-Rad), washed with 20 column volumes of buffer (50 mM Tris-HCl, pH 8.0, 150 mM NaCl, 0.1% digitonin), and eluted using 3C protease cleavage at 4°C overnight for hAC9 and b9AC5 and 2 hours for hAC5 and AC95. The eluted protein was concentrated to 1 ml and subjected to size-exclusion chromatography using a Superose-6 Increase 10/300 GL column. Peak fractions corresponding to hAC9, hAC5, b9AC5 or AC95 were concentrated and used for Cryo-EM grid preparation.

**Cryo-EM sample preparation and data collection**. For cryo–electron microscopy (cryo-EM) analysis, freshly purified full-length hAC9, or chimeric b9AC5 or AC95 in 0.1% digitonin was concentrated to 4.5 mg $mL^{-1}$ and incubated with a two-fold molar excess of G$\alpha$s in the presence or absence of 5 mM $MnCl_2$, 2 mM $MgCl_2$, 1 mM ATP$\alpha$S, and 0.5 mM forskolin. Aliquots (3.5 µL) were applied to glow-discharged Quantifoil R1.2/1.3 Cu grids, blotted for 3 s at blot force 20, and plunge-frozen in liquid ethane using a Mark IV Vitrobot at 100% humidity. Vitrified grids were stored in liquid nitrogen until data collection.

Cryo-EM data were collected at the ScopeM facility (ETH Zürich) on a 300 kV Titan Krios (FEI) equipped with a K3 direct electron detector (Gatan). Movies were recorded in super-resolution mode at a calibrated pixel size of 0.65 Å $pix^{-1}$, with a defocus range of −0.5 to −3.0 µm and fractionated into 40 frames. The total electron dose was 57 $e^-$ $Å^{-2}$ for the hAC9–G$\alpha$s dataset, 50 $e^-$ $Å^{-2}$ for b9AC5–G$\alpha$s dataset and 50 $e^-$ $Å^{-2}$ for both the hAC9–Gαs–ATP$\alpha$S–forskolin and AC95–Gαs–ATP$\alpha$S–forskolin datasets.

**Cryo-EM image processing**. Cryo-EM data processing was carried out using cryoSPARC (v4.0) [1]. Data processing workflows for hAC9-Gαs, hAC9–Gαs–ATP$\alpha$S–forskolin and AC95–Gαs–ATP$\alpha$S–forskolin are illustrated in Supplementary Figures 2-4. Large movie datasets recorded with a Titan Krios microscope (18,604 for hAC9-Gαs, 24,455 for hAC9–Gαs–ATP$\alpha$S–forskolin10524 for b9AC5 and 18,992 for AC95–Gαs–ATP$\alpha$S–forskolin). Movie frames were first aligned and corrected for beam-induced motion using Patch Motion. Contrast transfer function (CTF) parameters were estimated with Patch CTF, and micrographs of insufficient quality were excluded from further analysis. Particles were initially picked using template-based autopicking, guided by templates generated from a subset of manually picked particles. The resulting refined particle sets (6,453,541 for hAC9-G$\alpha$s, 6,856,880 for hAC9–Gαs–ATP$\alpha$S–forskolin, 3,103,570 for b9AC5 and 5,689,514 for AC95–Gαs–ATP$\alpha$S–forskolin). Multiple rounds of 2D classification were then performed to remove false positives and poorly aligned particles. Selected particle classes were subjected to ab initio reconstruction followed by non-uniform 3D refinement to generate an initial 3D map. For b9AC5, the resulting initial reconstruction was limited to low resolution, likely reflecting insufficient particle quality or dataset heterogeneity; therefore, processing was discontinued at this stage. To capture conformational heterogeneity, 3D variability analysis was applied, allowing the identification of distinct structural states [2]. These particle sets were refined using iterative cycles of additional rounds of 3D classification, non-uniform refinement, and per-particle CTF refinement [3]. For regions of particular interest, the catalytic domains, a focused mask was applied to perform local refinement, improving map quality in these regions.

The final density maps for the catalytic domain of AC bound to G$\alpha$s, generated with C1 symmetry applied, reached resolutions of 3.46 Å (hAC9-G$\alpha$s), 3.19 Å (hAC9–G$\alpha$s–ATP$\alpha$S–forskolin, state1), 3.57 Å (hAC9–G$\alpha$s–ATP$\alpha$S–Forskolin, state 2), and 3.55 Å (AC95–G$\alpha$s–ATP$\alpha$S–forskolin) as determined by Fourier shell correlation at the 0.143 threshold between independently refined half-maps. Finally, the resulting maps were subjected to automated post-processing and sharpening with DeepEMhancer (v0.16) [4], which improved side-chain resolution and ligand density, thereby yielding reconstructions suitable for reliable model building and structural interpretation.

**Model building and refinement**. AlphaFold 3–predicted models of hAC9 and AC95 in complex with G$\alpha$s were docked into the cryo-EM reconstructions using UCSF ChimeraX (v1.10.1) [5]. The models were iteratively rebuilt through cycles of manual adjustment in Coot (v0.9.8) [6] and real-space refinement in phenix.real_space_refine (Phenix v2.0) [7]. AlphaFold 3 predictions were used to guide backbone rebuilding in regions of lower local resolution [8]. Refinement was performed with restraints on secondary structure elements, forskolin, and ATP$\alpha$S. After each refinement cycle, model geometry and map fit were assessed, followed by targeted manual corrections. This process was repeated until convergence was achieved. Model quality was evaluated using MolProbity (v4.5.2) [9].

**Adenylyl cyclase activity assays**. The cAMP accumulation assay was performed using purified hAC9, hAC5, and AC95. Reactions were assembled with hAC9, hAC5, or AC95 alone or in complex with G$\alpha$s at a 1:2 molar ratio, in the presence of 0.5 mM allosteric or catalytic modulators. The assay was initiated by mixing 100 µL of protein solution with 100 µL of reaction buffer containing 4 mM $MgCl_2$, 10 mM $MnCl_2$, 0.2 mM ATP, and 20 nM [$^3$H]-ATP. Reactions were incubated at 30 °C for 30 min, then quenched by adding 20 µL of 2.2 M HCl, followed by heating at 95 °C for 4 min. Samples were rapidly cooled on ice and applied to gravity-flow columns containing 1.3 g of aluminum oxide. cAMP was selectively eluted with 4 mL of 0.1 M ammonium acetate. The eluate was mixed with 12 mL of scintillation cocktail, and radioactivity was quantified using a Packard 2250 CA Tri-Carb liquid scintillation counter.

### SUPPLEMENTARY FIGURES

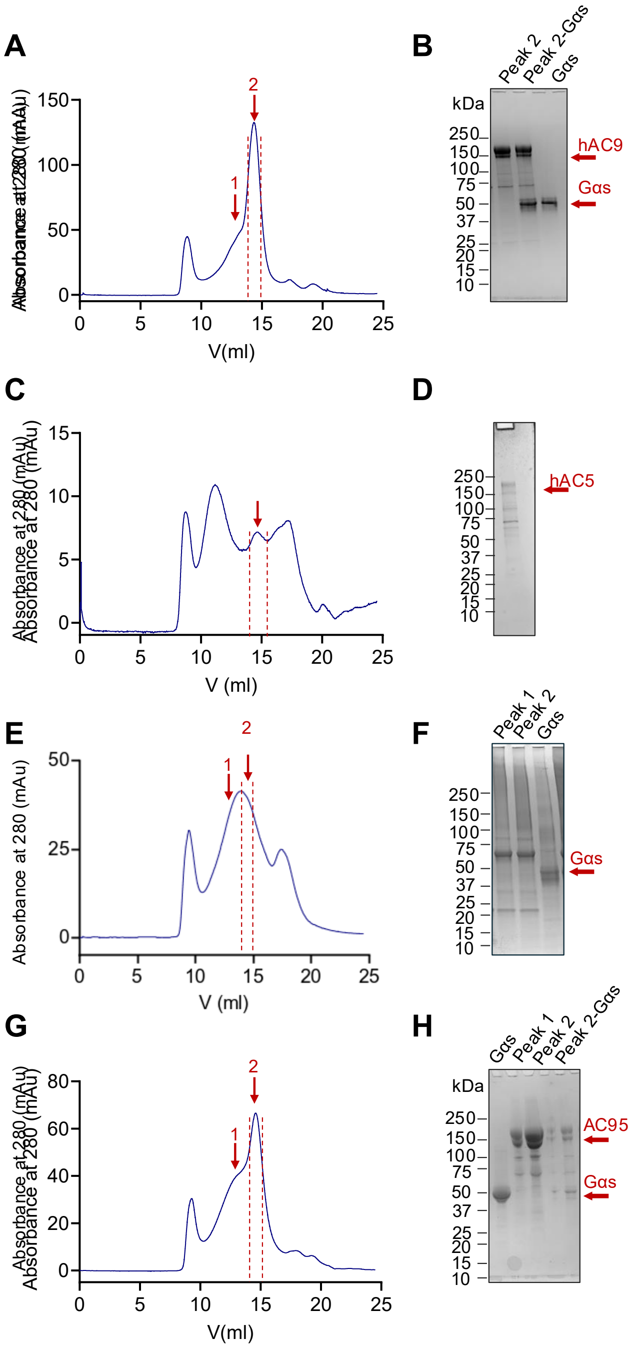


**Figure S1: Purification of hAC9, hAC5, b9AC5 and AC95. A.** Size-exclusion chromatography (SEC) profile of purified hAC9. **B.** SDS–PAGE analysis of purified hAC9 visualized by Coomassie staining. **C.** Size-exclusion chromatography (SEC) profile of purified hAC5. **D.** SDS–PAGE analysis of purified hAC5 visualized by Coomassie staining. **E.** Size-exclusion chromatography (SEC) profile of purified b9AC5. **F.** SDS–PAGE analysis of purified b9AC5 visualized by Coomassie staining. **G.** Size-exclusion chromatography (SEC) profile of purified AC95. **H.** SDS–PAGE analysis of purified AC95 visualized by Coomassie staining.

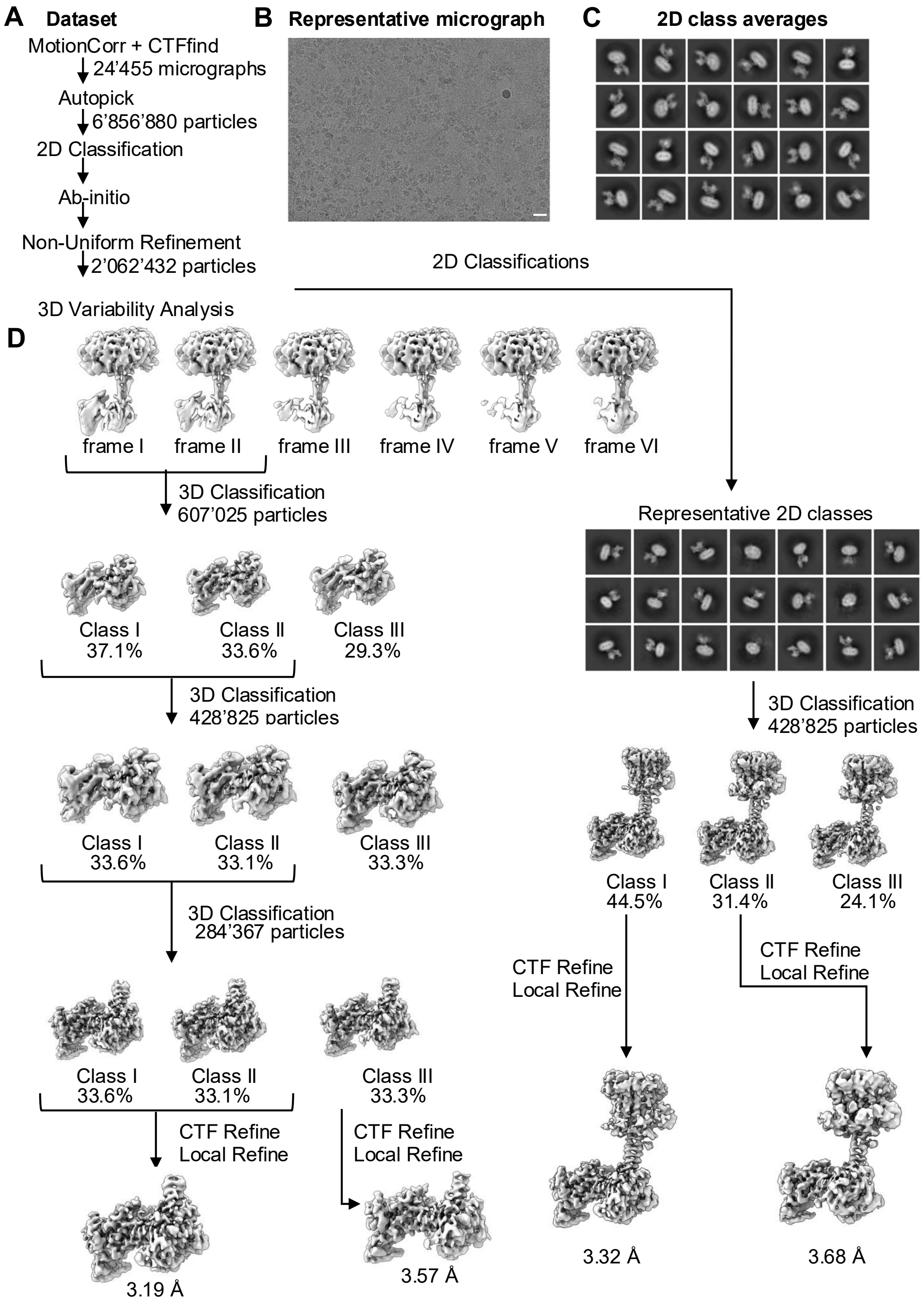


**Figure S2: Cryo-EM workflow for hAC9-Gαs-ATPαS-forskolin. A.** Cryo-EM data processing workflow for the human hAC9–Gαs–ATPαS–forskolin complex. **B.** Representative cryo-EM micrograph of the hAC9–Gαs–ATPαS–forskolin sample (scale bar, 20 nm). **C.** Selected 2D class averages (box size, 400 Å). **D.** Following 2D classification, 3D variability analysis, 3D classification, and local refinement of the catalytic domain, two conformations were resolved: (i) occluded state, state 1, at 3.19

Å, and (ii) ATPαS-bound state, state 2, at 3.57 Å resolution. The 3D reconstructions for the full-length proteins are at 3.32 Å and 3.68 Å resolution, respectively.

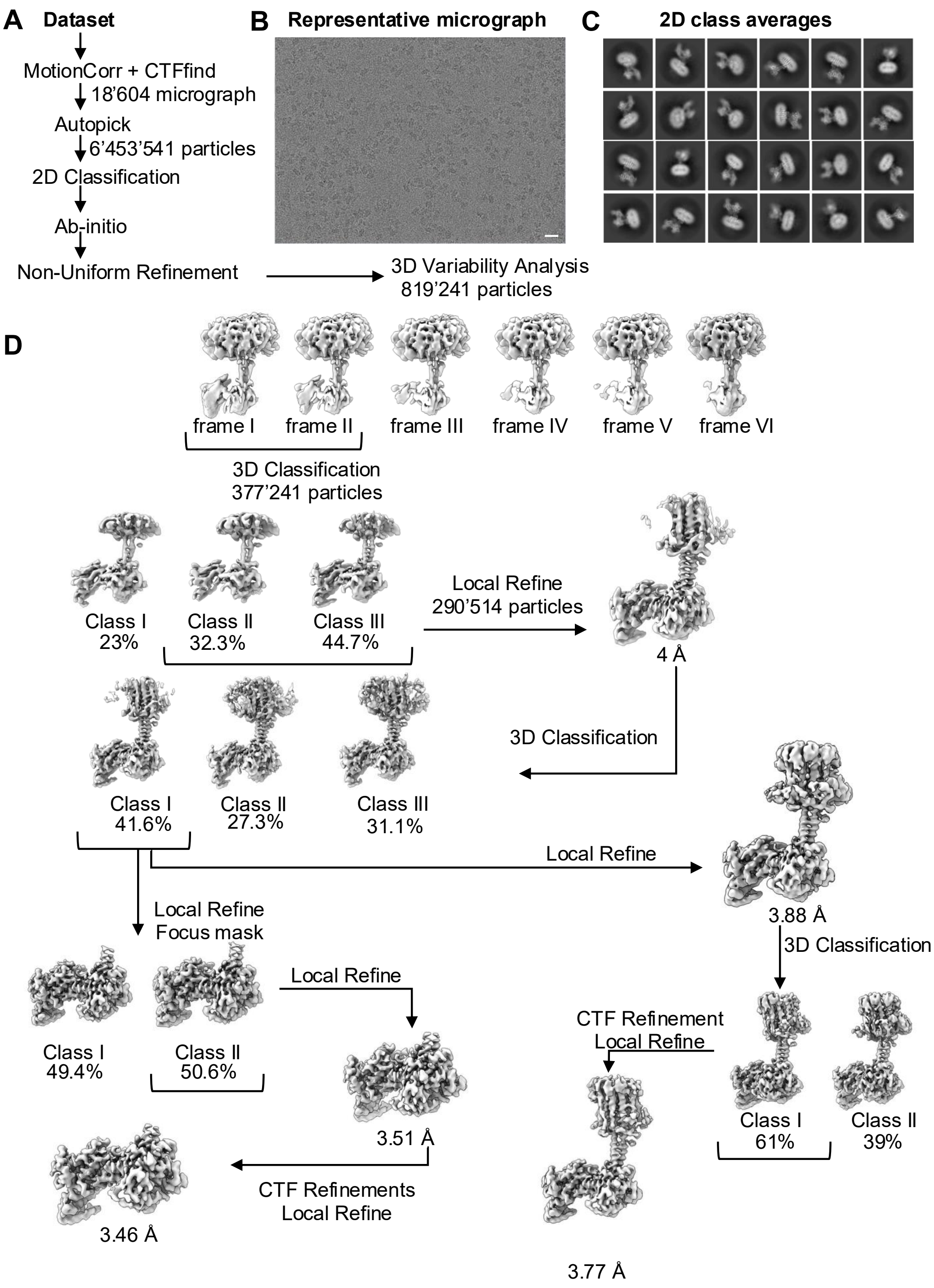


**Figure S3: Cryo-EM workflow of hAC9-Gαs. A.** Cryo-EM data processing workflow for the human hAC9–Gαs complex. **B.** Representative cryo-EM micrograph of the hAC9–Gαs sample (scale bar, 20 nm). **C.** Selected 2D class averages (box size, 400 Å). **D.** Following 2D classification, 3D variability analysis, 3D classification, and local refinement of the catalytic core and full-length AC9, the final reconstructions reached 3.77 Å for the full-length complex and 3.46 Å for the locally refined map.

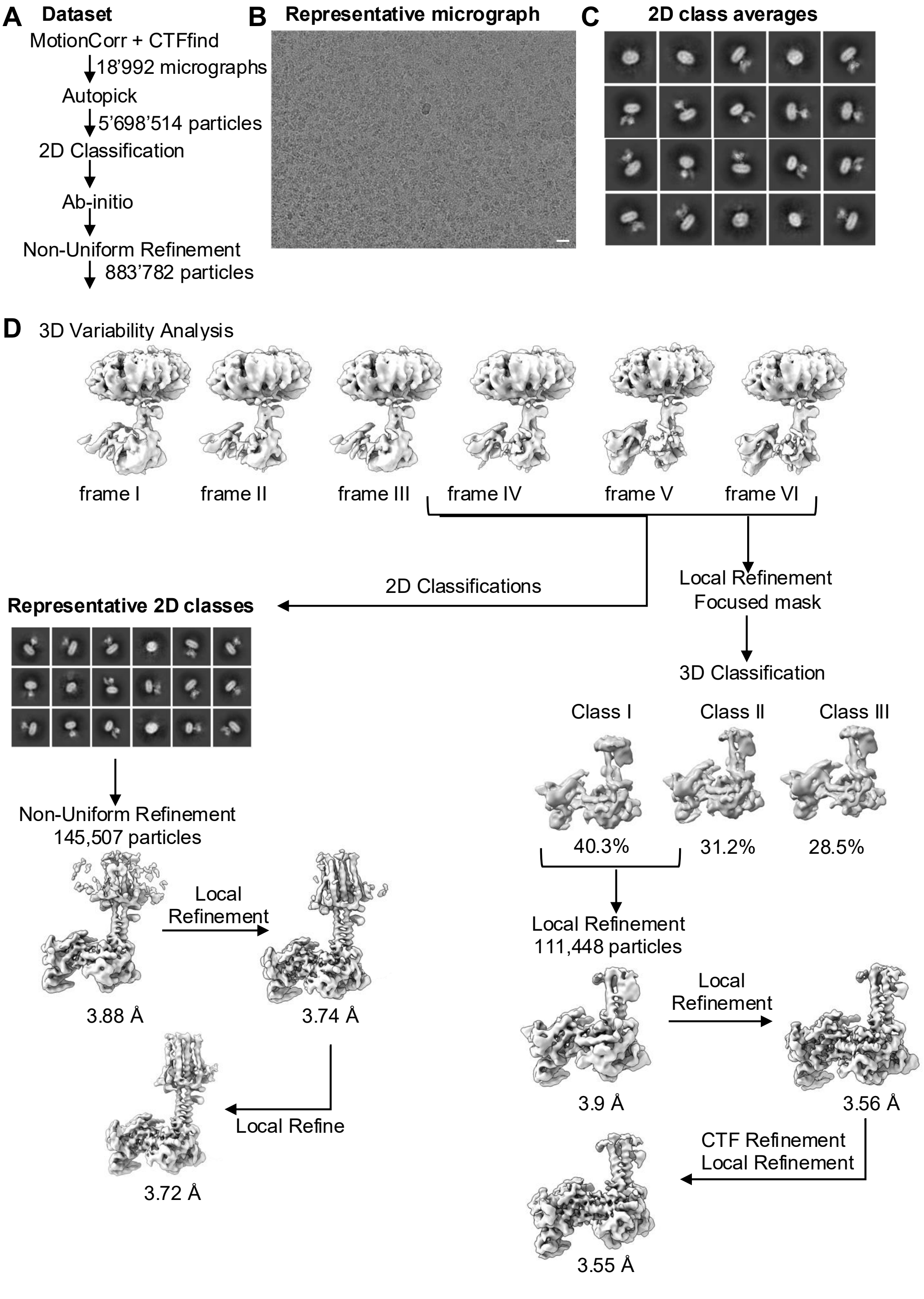


**Figure S4: Cryo-EM workflow of AC95-Gαs-ATPαS-forskolin. A.** Cryo-EM data processing workflow for the AC95–Gαs complex. **B.** Representative cryo-EM micrograph of the AC95–Gαs–ATPαS–forskolin sample (scale bar, 20 nm). **C.** Selected 2D class averages (box size, 400 Å). **D.** Following 2D classification, 3D variability analysis, 3D classification, and local refinement of the catalytic core and full-length AC95, the final reconstructions reached 3.72 Å for the full-length complex and 3.55 Å for the locally refined map.

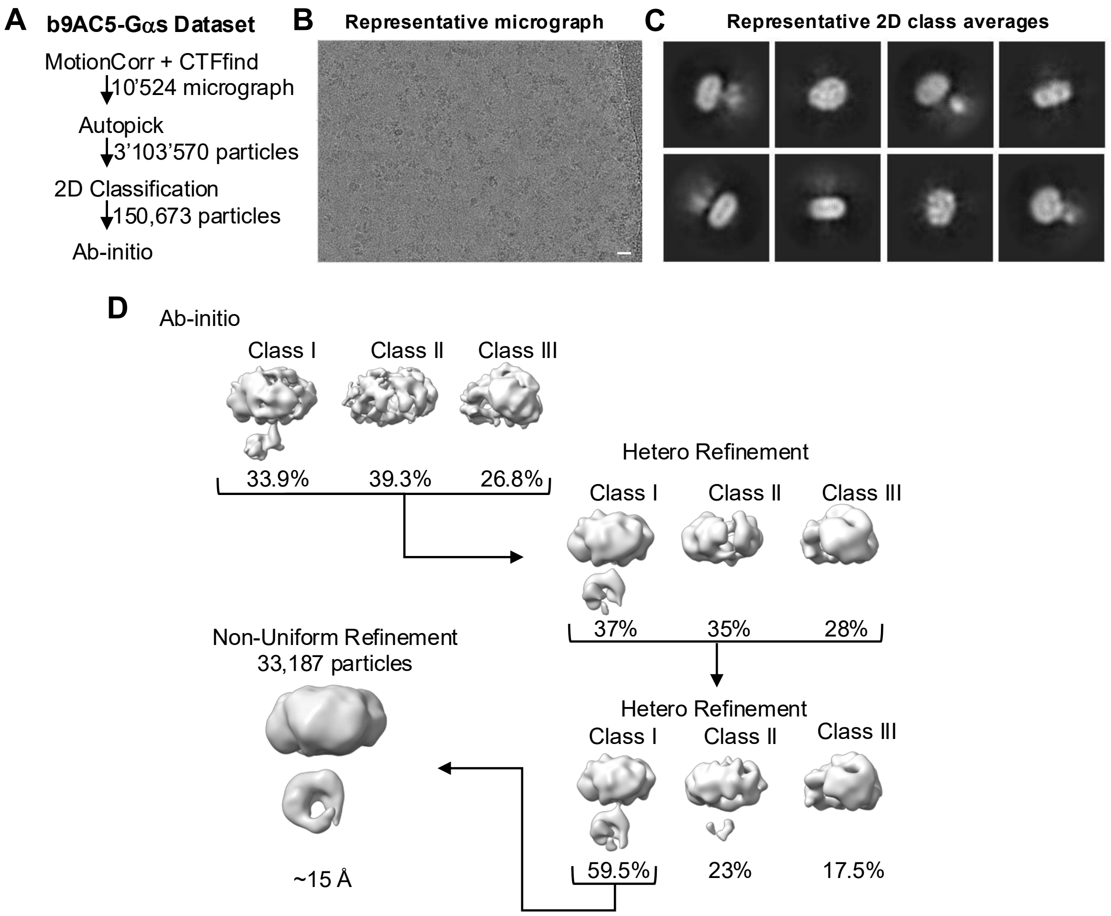


**Figure S5: Cryo-EM workflow of b9AC5-Gαs. A.** Cryo-EM data processing workflow for the b9AC5–Gαs complex. **B.** Representative cryo-EM micrograph of the b9AC5–Gαs complex (scale bar, 20 nm). **C.** Selected 2D class averages (box size, 400 Å). **D.** Following 2D classification, ab-initio, heterogenous refinement and non-uniform refinement, a low-resolution reconstruction of the b9AC5–Gαs complex was obtained (~15 Å resolution), reflecting the limited utility of the b9AC5 construct for structural studies.

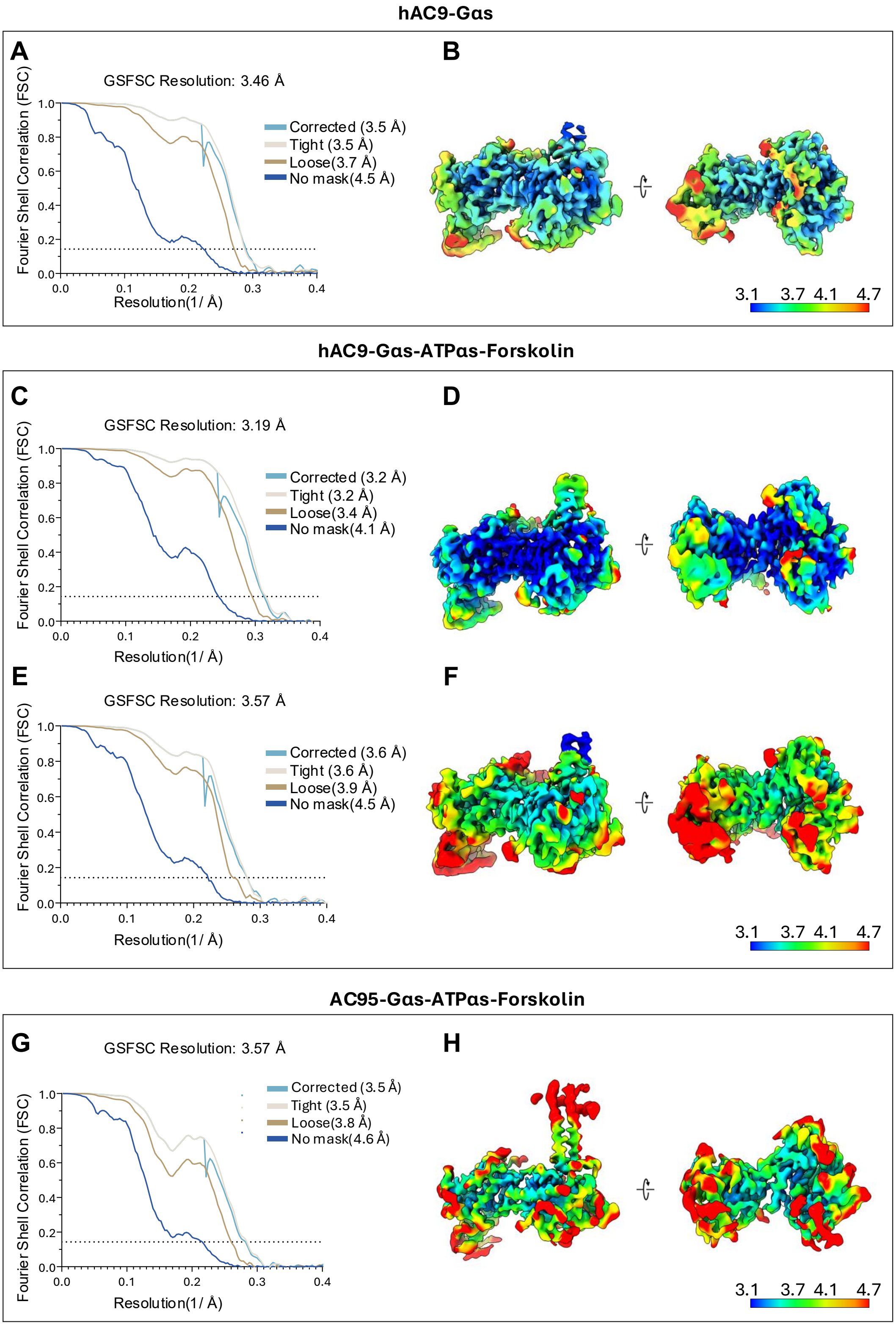


**Figure S6: Cryo-EM reconstructions and resolution assessment of catalytic domains of hAC9-Gαs, hAC9-Gαs-ATPαS-forskolin, and AC95-Gαs-ATPαS-forskolin samples. A-B.** Gold-standard Fourier shell correlation (GSFSC) curves (A) and local resolution map of hAC9–

Gαs 3D reconstruction, colored by local resolution (3.1–4.7 Å). **C-D.** Same as A-B, for hAC9–Gαs occluded state. **E-F.** Same as A-B, for hAC9–Gαs–ATPαS–forskolin (ATPαs-bound state). **G-H.** Same as A-B, for AC95–Gαs–ATPαs–forskolin.

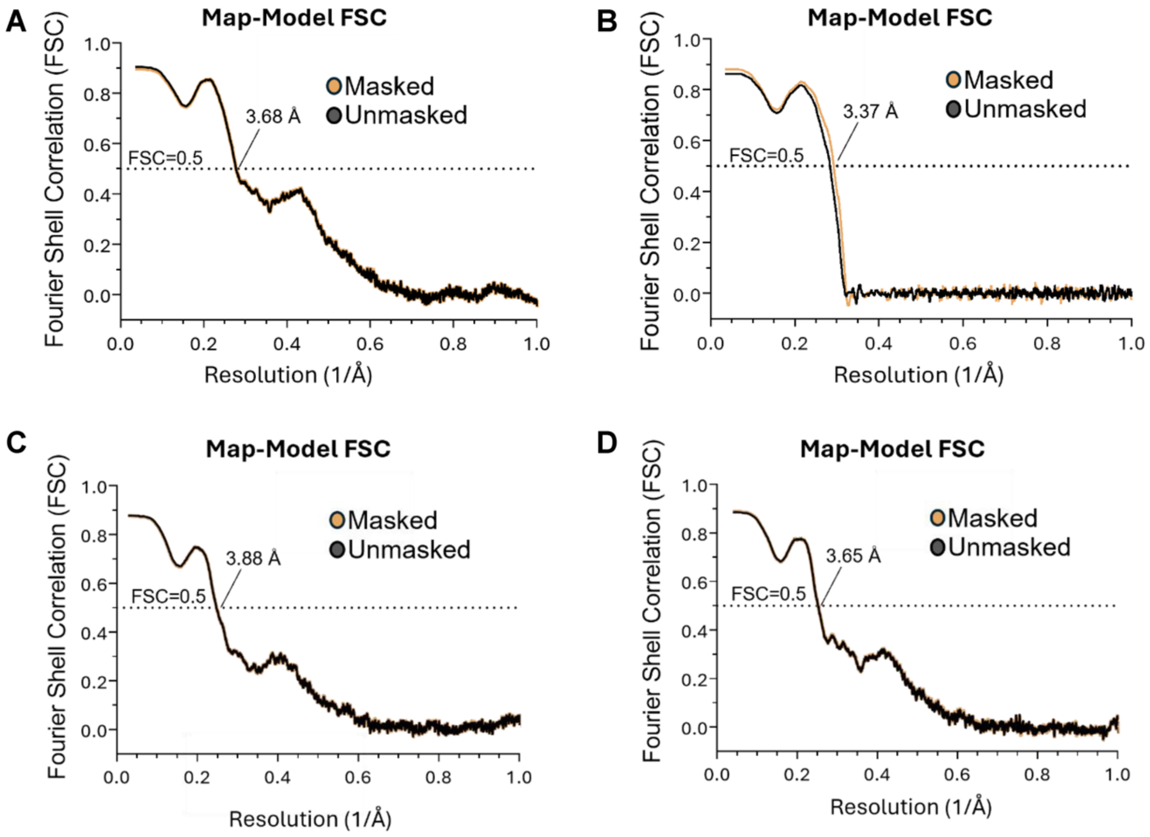


**Figure S7: Map-model FSC. A.** Map–model Fourier shell correlation (FSC) curve for the hAC9–Gαs complex, showing an FSC = 0.5 cutoff at 3.68 Å resolution. **B.** Map–model Fourier shell correlation (FSC) curve for the hAC9–Gαs-ATPαS-forskolin complex in occluded state, showing an FSC = 0.5 cutoff at 3.37 Å resolution. **C.** Map–model Fourier shell correlation (FSC) curve for the hAC9–Gαs-ATPαS-forskolin complex in ATPαS-bound state, showing an FSC = 0.5 cutoff at 3.88 Å resolution. **D.** Map–model Fourier shell correlation (FSC) curve for the AC95–Gαs-ATPαS-forskolin complex, showing an FSC = 0.5 cutoff at 3.65 Å resolution.

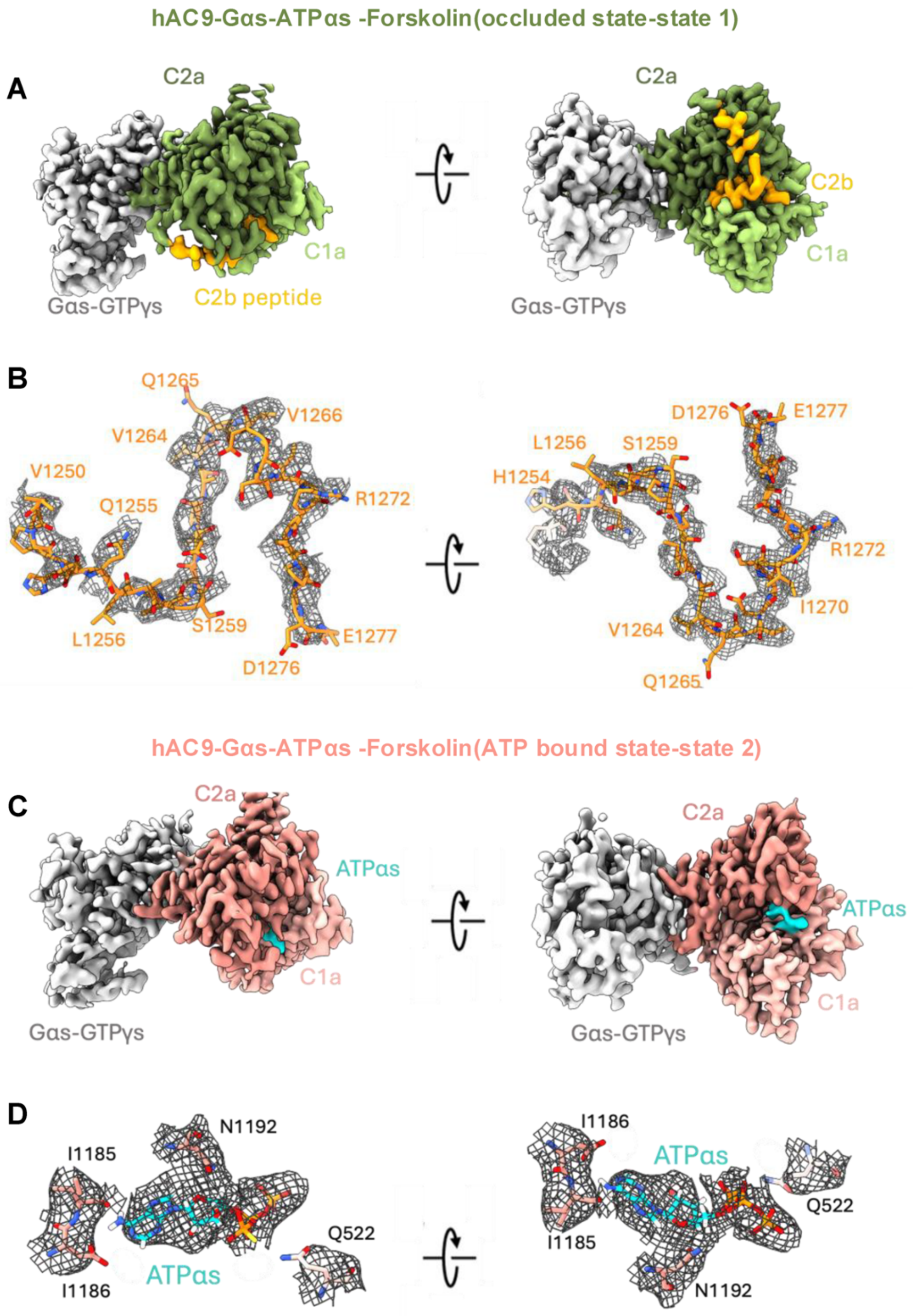


**Figure S8: hAC9 catalytic core with C2b peptide or ATPαS. A.** Focused view of the catalytic domain of the hAC9–Gαs-ATPαS-forskolin complex in the occluded state. **B.** C2b peptide residues with surrounding cryo-EM density shown as a mesh. **C.** Catalytic domain of hAC9–Gαs-ATPαs-Forskolin bound to ATPαS; Forskolin was present in the sample but not observed in the density. **D.** Residues within ~3.5 Å of ATPαs, with cryo-EM density displayed as a mesh.

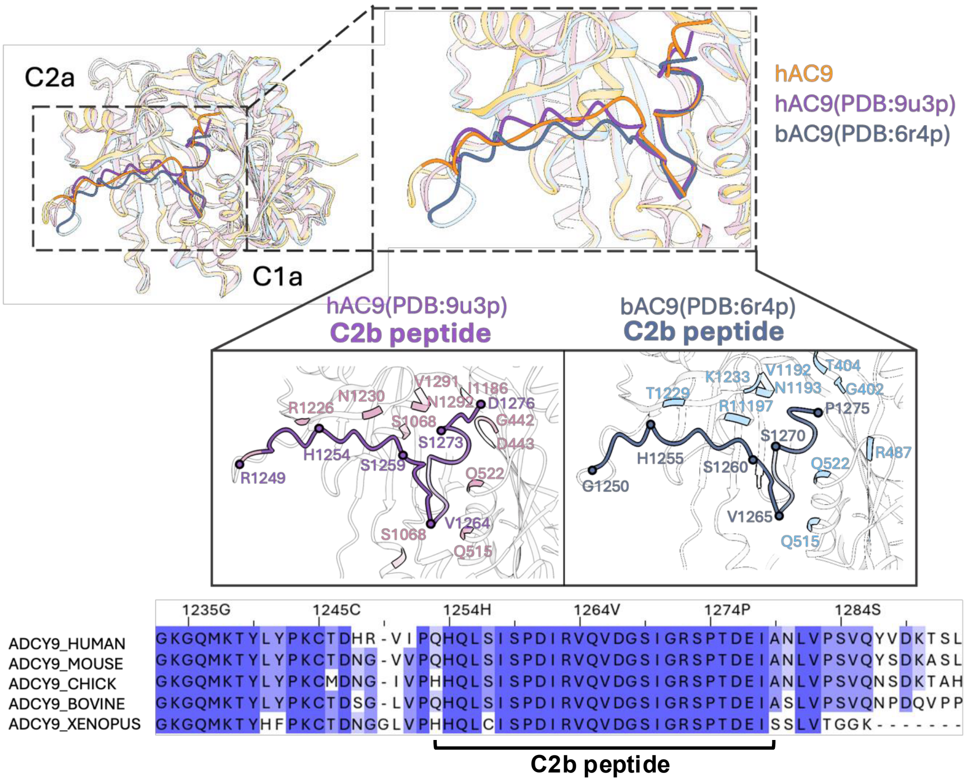


**Figure S9: Sequence conservation of the AC9 C2b peptide across species.** Multiple sequence alignment of the C2b peptide from AC9 across different organisms, highlighting conserved residues and motifs that suggest evolutionary conservation of its regulatory role**.**

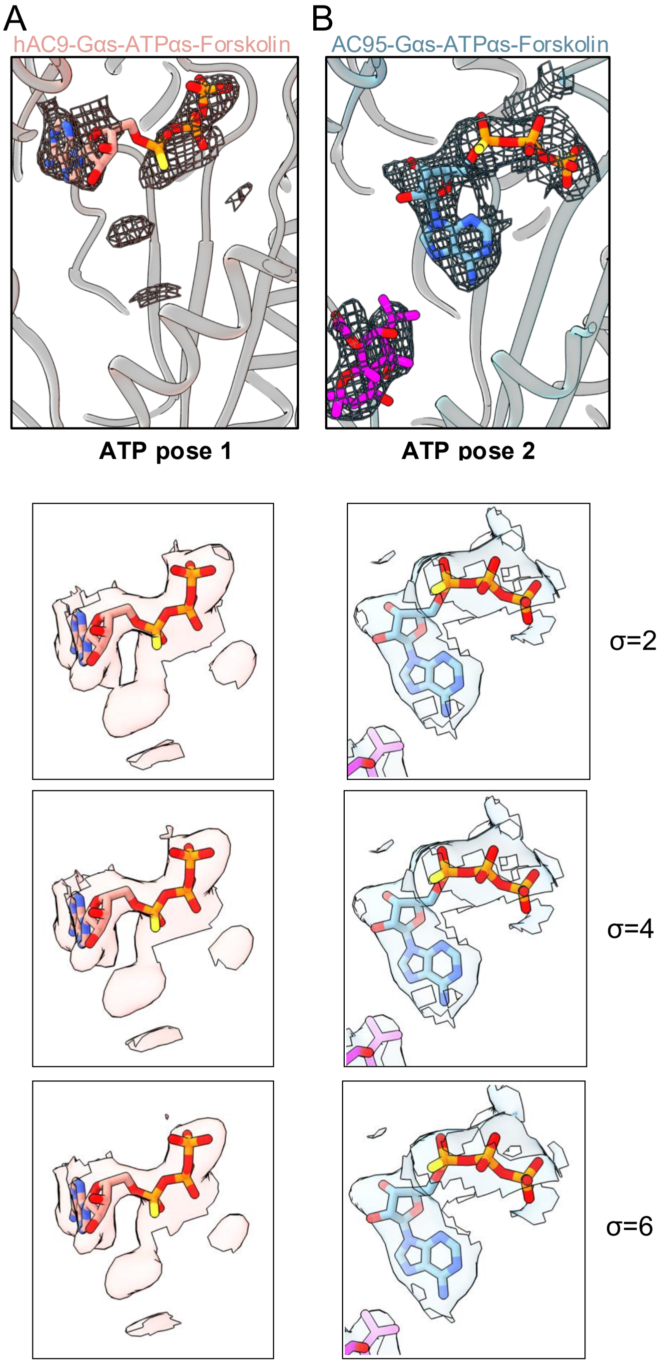


**Figure S10: Distinct ATPαS poses in hAC9 and AC95. A.** The observed ATPαS pose in hAC9-Gαs-ATPαS-forskolin sample, ATPαS pose 1. **B.** The observed ATPαS pose in AC95-Gαs-ATPαS-forskolin sample, ATPαS pose 2, in presence of forskolin. The density for ATPαS is shown at different σ levels for both hAC9 and AC95.

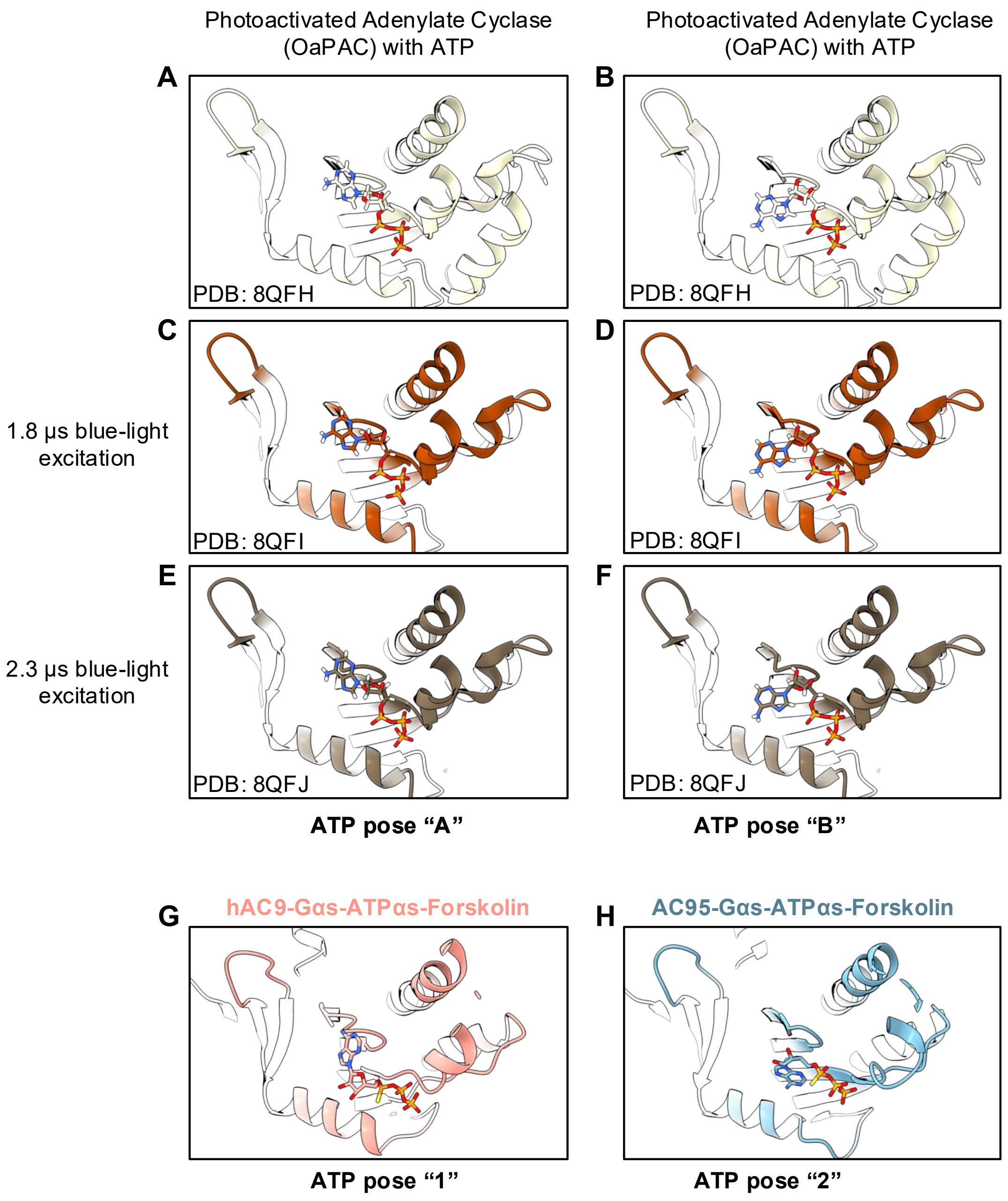


**Figure S11: Comparison of nucleotide-bound states. A-B.** A distinct ATP pose observed in the photoactivated AC, OaPAC [10]. **C-D.** Similar ATP poses observed upon blue light activation for 1.8 μs [10]. **E-F.** Similar ATP poses observed upon blue light activation for 2.3 μs [10]. **G.** ATP pose 1 observed in hAC9. **H.** ATP pose 2 observed in AC95.

**Supplementary Table S1. Cryo-EM analysis and statistics**

| Data Collection | | | | |
|---|---|---|---|---|
| Sample | AC9-Gαs | AC9-Gαs-ATPαS-forskolin | | AC95-Gαs-ATPαS-forskolin |
| Instrument | FEI Titan Krios / Gatan K3 Summit | | | |
| Voltage (kV) | 300 | | | |
| Electron Dose (e-/Å2) | 59 | 58 | | 50 |
| Defocus range (µm) | -0.5 to –3 | | | |
| Pixel size (Å) | 0.65 | | | |
| Number of particles | 57020 | State 1 occluded 190515 | State 2 ATPαS-bound 94694 | 111448 |
| FSC threshold 0.143 | 3.4 | 3.3 | 3.57 | 3.55 |
| **Refinement** | | | | |
| Model resolution FSC threshold 0.5 | 3.68 | 3.37 | 3.88 | 3.65 |
| Map CC | 0.76 | 0.73 | 0.68 | 0.72 |
| **Model composition** | | | | |
| Protein residues/ligands/water | 792/0/0 | 803/0/0 | 755/1/0 | 752/2/0 |
| Bond length r.m.s.d (Å) | 0.003 | 0.002 | 0.014 | 0.015 |
| Bond angle r.m.s.d (°) | 0.532 | 0.511 | 1.422 | 1.576 |
| **Validation** | | | | |
| MolProbity score | 1.66 | 1.68 | 1.89 | 2.13 |
| Clash score | 7.38 | 4.79 | 11.59 | 17.92 |
| Rotamer outlier (%) | 0.3 | 2.48 | 0.62 | 0.17 |
| Ramachandran plot | | | | |
| Favoured (%) | 96.33 | 97.01 | 95.5 | 94.41 |
| Allowed (%) | 3.67 | 2.99 | 4.5 | 5.59 |
| Disallowed (%) | 0.00 | 0.00 | 0.00 | 0.00 |
